# Engineering Defensin α-helix to produce high-affinity SARS-CoV-2 Spike protein binding ligands

**DOI:** 10.1101/2022.02.09.479781

**Authors:** Leonardo Antônio Fernandes, Anderson Albino Gomes, Maria de Lourdes Borba Magalhães, Partha Ray, Gustavo Felippe da Silva

**Affiliations:** Biochemistry Laboratory, Center of Agroveterinary Sciences, State University of Santa Catarina, Lages, Santa Catarina, 88520-000, Brazil; Division of Surgical Oncology, Department of Surgery, Moores Cancer Center, University of California - San Diego Health, La Jolla, CA 92093, USA

**Keywords:** COVID19, Spike-protein, ACE2, Defensin, Peptidomimetics, SARS-CoV-2 Diagnostics

## Abstract

The binding of Severe Acute Respiratory Syndrome Coronavirus 2 (SARS-CoV-2) Spike protein to the Angiotensin-Converting Enzyme 2 (ACE2) receptor expressed on the host cells is a critical initial step for viral infection. This interaction is blocked through competitive inhibition by soluble ACE2 protein. Therefore, developing high-affinity and cost-effective ACE2 peptidomimetic ligands that disrupt this protein-protein interaction is a promising strategy for viral diagnostics and therapy. We employed human and plant defensins, a class of small and highly stable proteins, and engineered the amino acid residues on its conformationally constrained alpha-helices to mimic the critical residues on the ACE2 helix 1 that interacts with the Spike-protein. The engineered proteins were soluble and purified to homogeneity with high yield from a bacterial expression system. The proteins demonstrated exceptional thermostability, high-affinity binding to the Spike protein with dissociation constants in the low nanomolar range, and were used in a diagnostic assay that detected SARS-CoV-2 neutralizing antibodies. This work addresses the challenge of developing helical peptidomimetics by demonstrating that defensins provide promising scaffolds to engineer alpha-helices in a constrained form for designing high-affinity ligands.

**Broad audience statement:** The engineered proteins developed in this study are cost-effective and highly stable reagents for SARS-CoV-2 detection. These features may allow large-scale and cost-effective production of diagnostic tests to assist COVID-19 diagnostic and prevention.

## Introduction

Coronavirus Disease 2019 (COVID-19) caused by Severe Acute Respiratory Syndrome Coronavirus 2 (SARS-CoV-2) emerged as a pandemic in March 2020 (1, 2), resulting in over 300 million infections and more than 5.5 million deaths worldwide as of January 2022 (3). The emergence of highly transmissible variants and the possible occurrence of other pandemics calls for innovative protein engineering strategies to develop improved viral detection probes for rapid diagnostic followed by proper therapeutic interventions. The viral entry into human cells occurs when SARS-CoV-2 Spike glycoprotein (S) binds the human Angiotensin-Converting Enzyme 2 (ACE2) followed by its proteolytic cleavage by the cellular Trans-Membrane Protease Serine 2 (TMPRSS2) (4). Therefore, binding to the human ACE2 receptor is a critical initial step for SARS-CoV-2 infection, and targeting such Protein-Protein Interactions (PPIs) via competitive-binding might be a promising strategy for blocking viral entry into human cells (5). Accordingly, studies have shown that soluble ACE2 inhibits SARS-CoV-2 infection by blocking the interaction between the human receptor and the viral protein (6) (7).

Biochemical and structural studies have demonstrated that interactions with the Receptor-Binding Domain (RBD) of Spike SARS-CoV-2 mainly involve ACE2 N-terminal α-helix 1 (8), and such α-helical peptide may effectively mimic ACE2 and bind to the Spike protein. However, once a short peptide is isolated from the original protein, much of its binding ability is lost, as the peptide tends to adopt random-coil conformations, reducing the number of optimal binding conformations (9–12).

Therefore, the development of stabilized α-helical mimetic ligands is crucial for Spike protein binding, allowing SARS-CoV-2 detection, COVID-19 neutralizing antibody analysis, and therapeutic intervention design. For that reason, several attempts to produce stable and potent α-helical ACE2 peptidomimetics are underway (13–16).

The proteolytic instability of peptides is an additional factor that limits their utility as reagents in diagnostic and therapy since flexible peptides are also easy targets for proteases (17). In this regard, reduced conformational heterogeneity of stabilized helical peptides has been demonstrated to increase protease resistance (12).

Defensins are small (45-54 amino acid residues), cysteine-rich, cationic proteins found in animals, plants, and fungi that play crucial roles in the innate immune system by protecting against pathogens (18, 19). These proteins usually enclose three to five disulfide bonds that confer protease resistance (20), extreme temperature tolerance (21), and wide pH-range stability (22).

In plants, all defensins share a common topology that consists of one α-helix and three antiparallel β-strands stabilized by at least four disulfide bonds (23). However, in vertebrates, α-helices are present only in a subset of defensins called beta-type that contain a α-helix restricted to the carboxy-terminal region of these proteins (18, 24).

In this study, we hypothesized that the design of ACE2 α-helix peptidomimetics using defensin scaffolds could be advantageous given their stability and helix-constrained conformation. Therefore, considering the published crystal structure of the SARS-CoV-2 RBD-ACE2 complex (25), we selected the main residues located on the α-helix 1 of ACE2, contacting RBD, and introduced them into the α-helix of plant and human defensins to investigate the ability of the engineered proteins to bind the RBD of Spike protein from SARS-CoV-2.

The use of prokaryotic expressed proteins presents several advantages, including production cost and time efficiency compared to antibodies or eukaryotic proteins. The engineered proteins developed in this study were highly expressed in *Escherichia coli*, binding SARS-CoV-2 Spike protein with low nanomolar *K*_d_ values and presenting exceptional thermal stability. In addition, proteins were able to detect HEK-293/SARS-CoV-2 Spike expressing cell lines and detected the presence of SARS-CoV-2 neutralizing antibodies via competitive ELISA tests. In summary, we report a novel strategy to generate constrained α-helix SARS-CoV-2 binders stabilized by defensin scaffolds that are promising candidates for COVID-19 diagnostics and viral neutralization.

## Results

### Design of defensin-based ACE2 peptidomimetics

To produce a potentially low-immunogenic and stable ACE2 mimic we initially chose a human scaffold (defensin PDB1IRH). Analysis of the complex ACE2-RBD structure using the Pymol software reveals that most RBD contacts involve the surface located ACE2 helix 1 residues L29, D30, K31, N33, H34, E35, E37, D38, and L39. Therefore, we inserted residues D30, K31, H34, and E35 into the human defensin α-helix to produce the engineered protein named h-deface2 (Figure 1, Supplementary Data S1). The chosen defensin scaffold contains a α-helix stabilized by two disulfide bonds; therefore, we hypothesized they would provide a constrained and stable framework to display ACE2 residues responsible for Spike RBD binding.

**Fig. 1:**
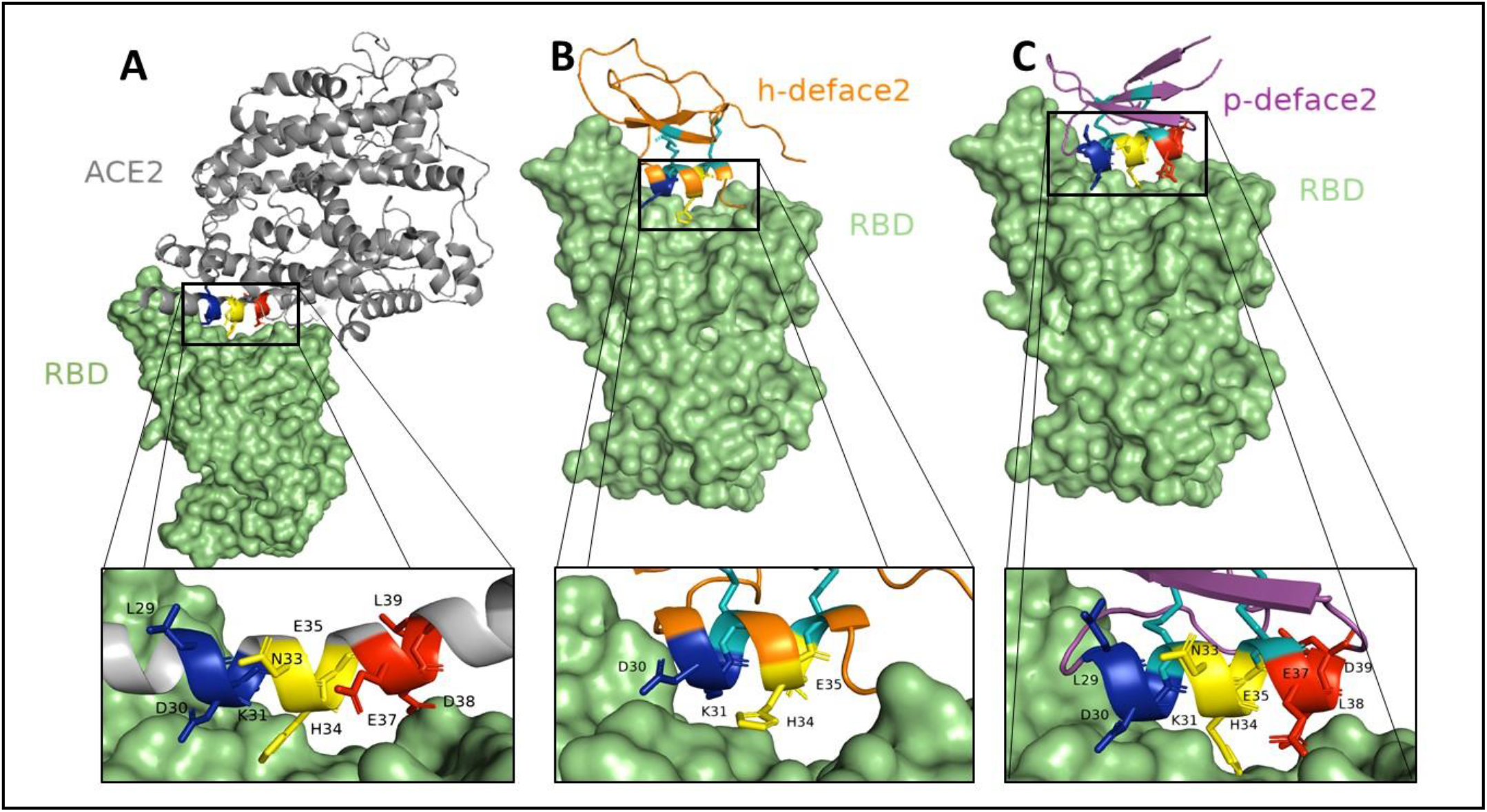
Schematic diagram of Defensin based-ACE2 peptidomimetics. Panel A: Three-dimensional structure of SARS CoV2 RBD/ACE2 complex highlighting the interactions between helix 1 and Spike RBD. Panel B: Hypothetical model of h-deface2 and RBD complex. The introduced h-deface2 α-helix residues were superimposed to ACE2 surface resides (D30, K31, H34, and E35) to create a hypothetical complex model. Panel C: Hypothetical model of p-deface2 and RBD complex. The p-deface2 a-helix residues (L29, D30, K31, N33, H34, E35, E37, D38, and L39) were superimposed to ACE2 surface resides to create the hypothetical complex model. The three-dimensional structure of defensin-based peptides were created using The Phyre2 web portal for protein modeling, prediction, and analysis (39). Disulfide bonds are shown in cyan.

Nonetheless, the human defensin contains a short α-helix that accommodates the insertion of only four α-helix ACE2 residues (Figure 1B). Therefore, we further selected a defensin from the tobacco plant (PDB 6MRY) that contained a more extended α-helical structure, which accommodated five additional ACE2 residues, intending to produce higher affinity binders (Figure 1C). Therefore, residues L29, D30, K31, N33, H34, E35, E37, D38, and L39 were introduced into the plant defensin α-helix, and the resulting protein was named p-deface2 (Figure 1). In addition, based on recent studies claiming increased binding upon ACE2 mutations E35K and K31N (26), we introduced such mutations on p-deface2 to produce a new version of the protein named p-deface2-MUT.

Engineered proteins were initially expressed as Trx-fusion peptides (Supplementary data S1) to assist disulfide bond formation, which led to soluble protein expression of pronounced yield (>50mg/L). To rule out the possibility of Trx-tag interference into the binding, Trx-free proteins were also expressed, purified, and tested accordingly. Both the Trx tagged and untagged proteins exhibited similar binding to the ECD of S-protein, demonstrating that the tag did not interfere with the binding (Supplementary data S2). However, heterologous expression of engineered peptides in the absence of Trx-tag resulted in a substantial yield decrease (< 5 mg/L).

### Determination of dissociation constant

Biolayer Interferometry (BLI) experiments were initially performed using biotinylated ACE2-peptidomimetics and SARS-CoV-2 S-protein to investigate the molecular interactions. Biotinylated h-deface2 and p-deface2 were immobilized on streptavidin-conjugated biosensors, and experiments were performed using a gradient of SARS-CoV-2 S-protein. Measured data indicates association and dissociation phases (Figure 2A and 2B), but curves did not follow a single exponential kinetic behavior. Therefore, kinetic analysis using either 1:1 or 1:2 binding models was not possible using this method, and so were the *K*_d_ calculations. We, therefore, calculated the apparent dissociation constants using dose-response curves from ELISA assays and the apparent *K*_d_ values were determined using nonlinear regression Michaelis-Menten curve fits (Figure 2C). Accordingly, calculated apparent *K*_d_ values were 54.4±11.3, 33.5±8.2, and 14.4±3.5 nM for h-deface2, p-deface2, and p-deface2-MUT, respectively (Table 1).

**Fig. 2:**
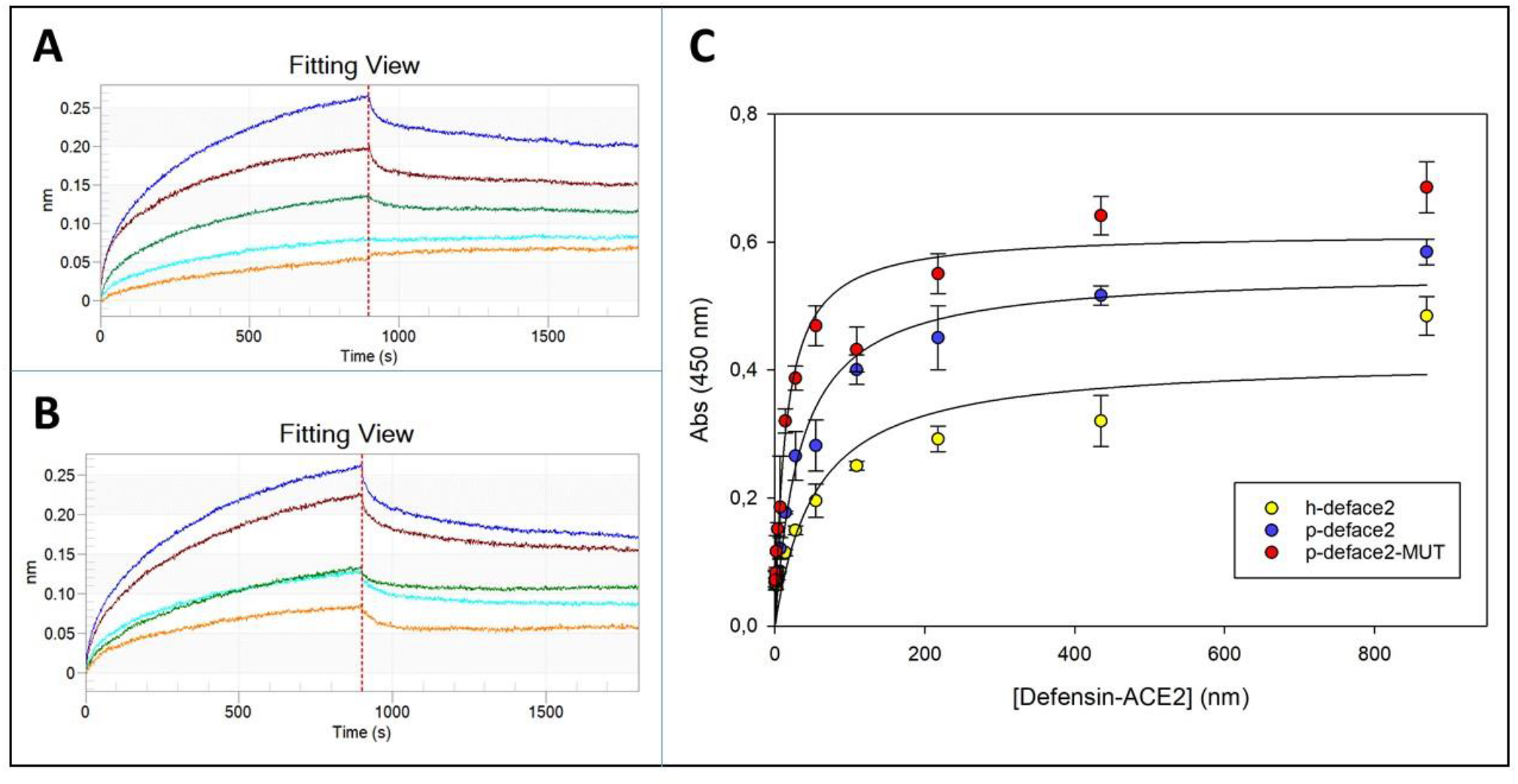
ACE2-Defensins binding to SARS-CoV-2 Spike protein. The binding of (A) h-deface2 and (B) p-deface2 to the RBD were monitored with BLI. Blue (40nM), red (30nM), green (20nM), cyan (10nM), and yellow (5nM) lines are measured data, and vertical red dotted lines indicate a transition between the association phase and dissociation phase. *K*_d_ could not be accurately estimated using BLI because curves did not follow a single exponential kinetic behavior. (C) h-deface2, p-deface2, and p-deface2MUT dose-response curves by direct. The apparent *K_d_* values were determined using the nonlinear regression Michaelis-Menten curve fit.

**Table 1:** The apparent dissociation constant (K_d_) values for Defensin-based ACE2 mimetic binding into SARS-CoV-2 Spike protein.

| ACE2-defensin | $K_d$ (nM) |
| --- | --- |
| h-deface2 | 54.4±11.3 |
| p-deface2 | 33.5±8.2 |
| p-deface-MUT | 14.4±3.5 |
| p-deface-ala | N/A |

Apparent $K_d$ values were determined using ELISA dose-response assays using variable ACE2 mimetic concentrations and nonlinear regression Michaelis- Menten curve fits. Measurements were performed in triplicates and errors correspond to standard deviation.

To test the binding specificity of the engineered ACE2-peptidomimetics, the amino-acid residues D30, K31, E35, D38, and L39 were mutated to alanine to produce p-deface2 ala mutant, and binding analysis was performed. We found that the alanine mutation disrupted binding to the SARS-CoV-2 S protein, thus reconfirming the importance of these residues for the binding interaction (Figure 3).

**Fig. 3:**
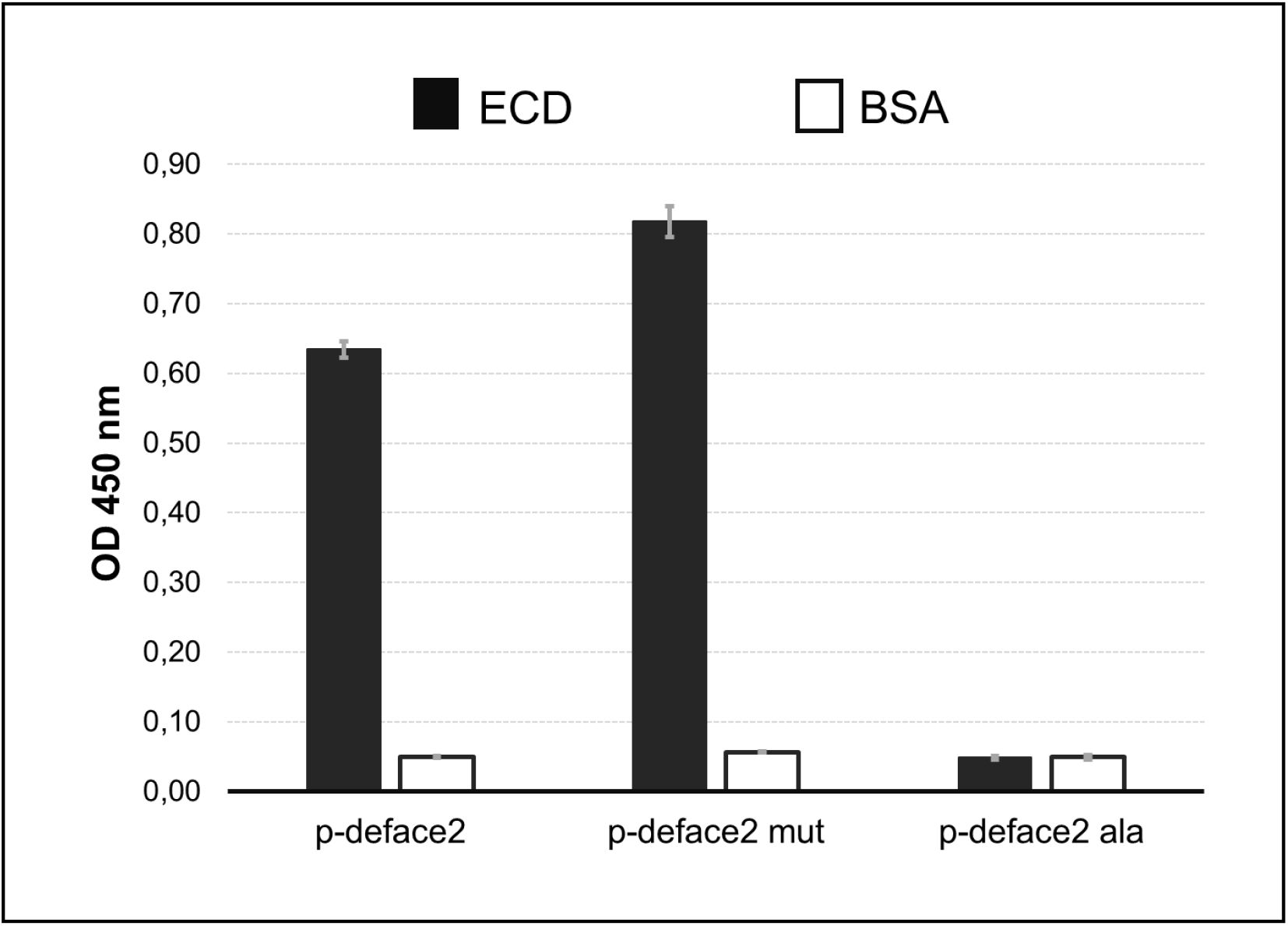
Mutation of key residues (D30, K31, E35, D38 and L39) to alanine disrupted SARS-CoV-2 S protein binding. 25 ng of SARS-CoV-2 S protein were immobilized into polystyrene plates and incubated with 100 ng of biotinylated p-deface2, p-deface-MUT, or p-deface-ala.

We also tested the thermostability of the engineered proteins. For this, we pre-incubated the purified proteins at specific temperatures and followed the binding using standard ELISA assay. We observed that all three proteins retained binding activity after heating (Figure 4). Additionally, the proteins retained their total activity upon lyophilization, followed by reconstitution (Supplementary data S3).

**Figure 4:**
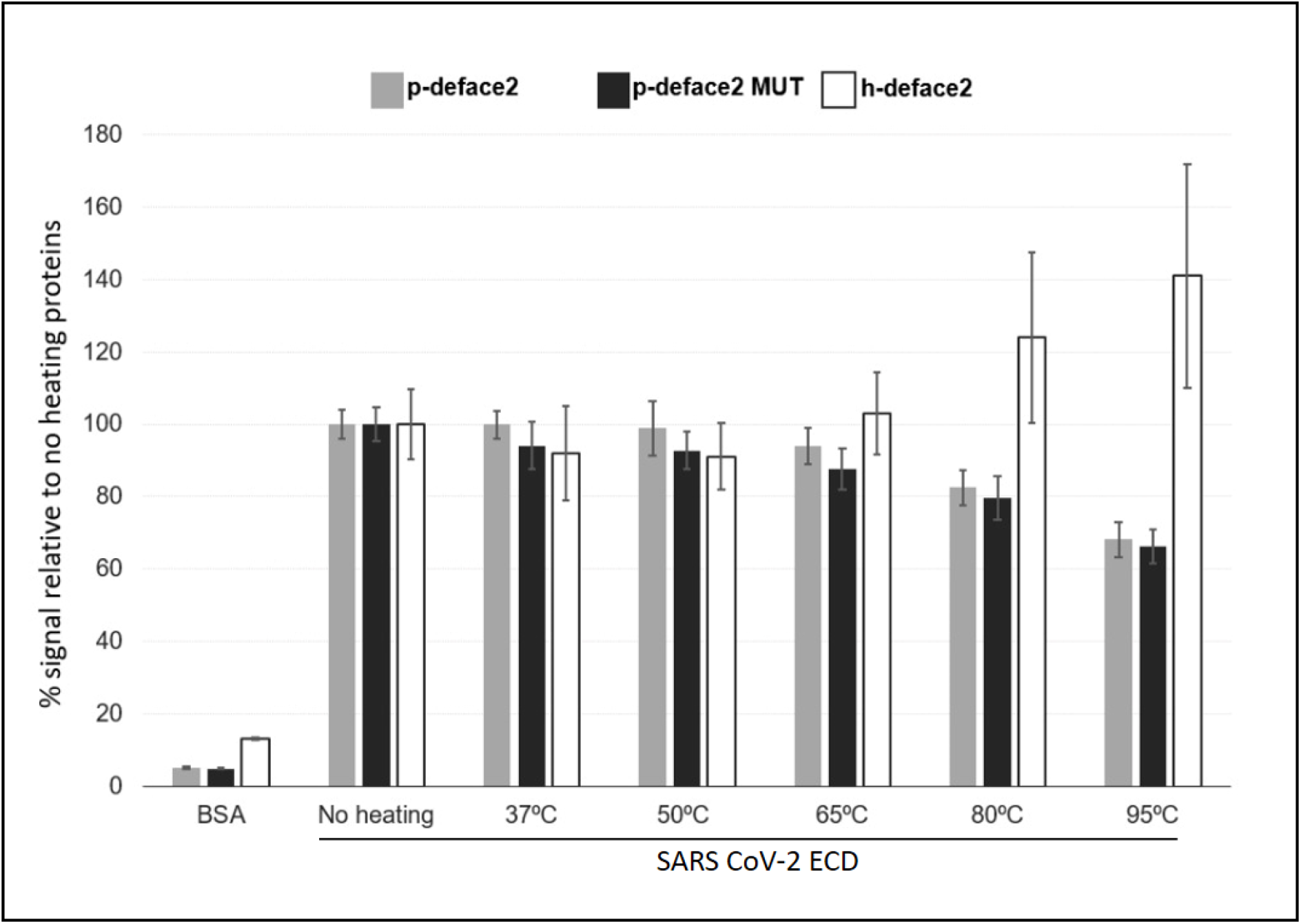
Defensin based-ACE2 peptidomimetics are thermostable. p-deface2 (gray bars), p-deface2-MUT (black bars), and h-deface2 (white bars) were incubated at temperatures ranging from 37 to 95 °C for 5 minutes, and binding against SARS-CoV-2 Spike protein was tested. 50 ng of ECD SARS-CoV-2 Spike was immobilized onto polystyrene ELISA plates following incubation with Defensin-ACE2 previously incubated at different temperatures.

### Diagnostic application of the ACE2-peptidomimetics

Next, we investigated the potential application of ACE2-peptidomimetics in a diagnostic platform for detecting SARS-CoV2 neutralizing antibodies. For this, we used a Brazilian Health Regulatory Agency (ANVISA)-approved neutralizing antibody test. The assay is based on the principle of competitive ELISA. Briefly, the biotinylated-ACE2 protein is immobilized on the streptavidin-coated microplates that can bind HRP-coupled SARS-CoV-2 Spike protein and generate a signal in the presence of HRP-substrate. However, in the presence of SARS-CoV2 anti-S-protein binding antibodies (neutralizing sera), the availability of HRP-coupled SARS-CoV-2 Spike protein to bind the ACE2 protein decreases, thus resulting in a decrease of the measured signal. We modified this assay and replaced the biotinylated-ACE2 with the biotinylated p-deface2 protein (Figure 5). This was followed by adding HRP-coupled SARS-CoV-2 Spike protein and the test (neutralizing) or control (non-neutralizing) human serum provided with the kit. The presence of neutralizing antibodies disrupted the interaction between p-deface2 and HRP-coupled SARS-CoV-2 Spike protein, thus decreasing the measured signal compared to the control non-neutralizing sera, indicating that biotinylated p-deface2 could be applied in a diagnostic assay platform to screen between positive and negative SARS-CoV-2 neutralizing serum (Figure 5).

**Fig. 5:**
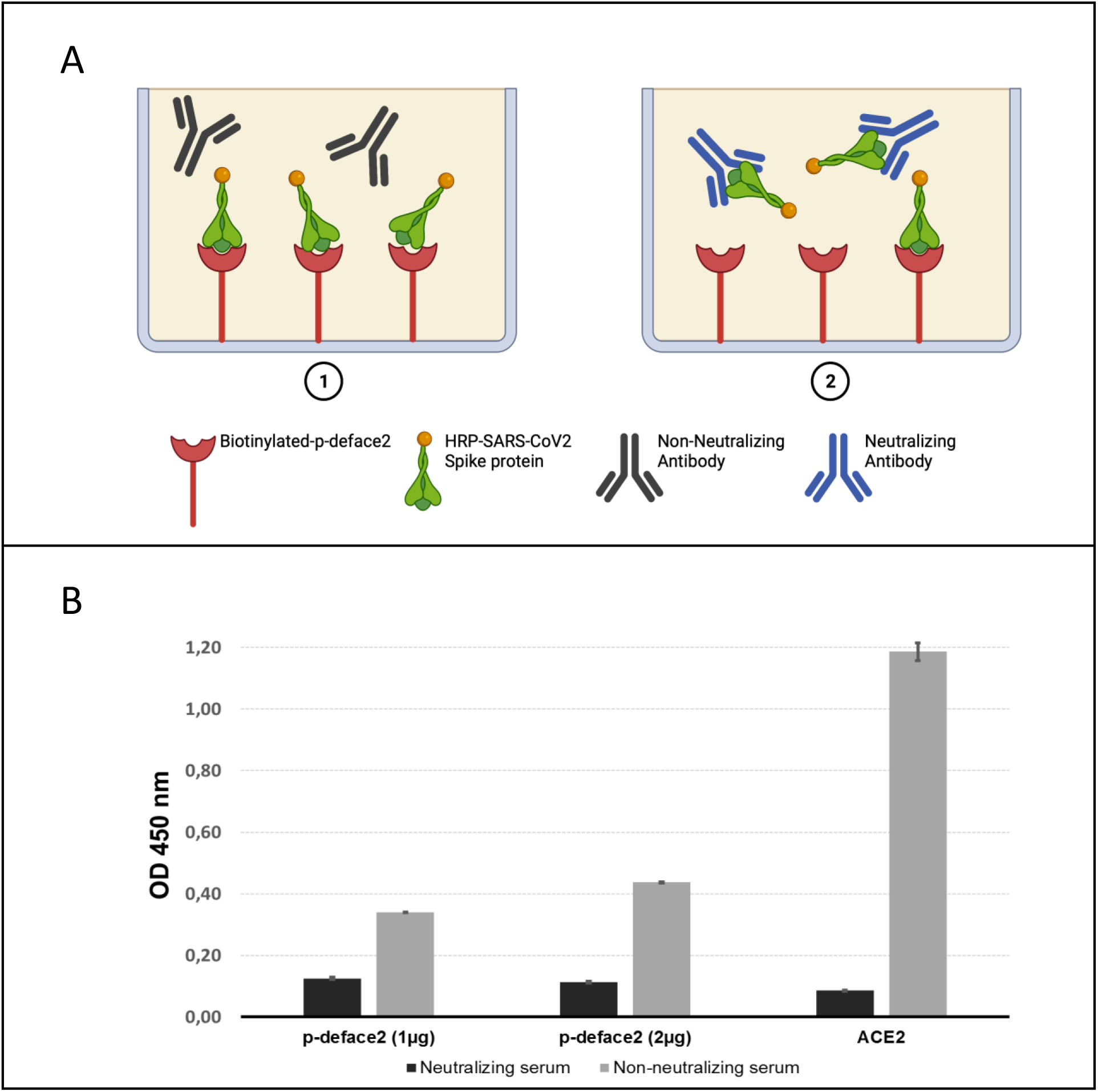
Detection of the presence of neutralizing antibodies using Defensin based-ACE2 peptidomimetics. Panel A: Schematic diagram of competitive ELISA to detect SARS-CoV-2 neutralizing antibody. In the presence of: (1) Non-neutralizing antibodies the HRP-coupled Spike protein binds to the immobilized p-deface-2. (2) Neutralizing antibodies block the HRP-coupled Spike and p-deface-2 interaction, resulting in decrease of signal in presence of HRP-substrate. The figure was created with BioRender.com. Panel B: ELISA assay. Biotinylated ACE2 or p-deface2 were immobilized into streptavidin-coated ELISA plates, followed by the addition HRP-coupled Spike protein in the presence of human serum. After incubation, wells were washed and incubated with TMB for 15 minutes.

### ACE2-peptidomimetics binding to cells expressing SARS-CoV2 Spike proteins

Finally, we wanted to determine if h-deface2 and p-deface2 could bind the S-protein expressed on the cell surface that might mimic the architecture of the trimeric-S-protein displayed on the SARS-CoV2 surface. Transgenic HEK293 cell lines stably transfected with (1) Wild-type (Wuhan-Hu-1) or (2) Delta-variant (B.1.617.2) S-protein expressing plasmids were used for this assay. First, we performed Flow-cytometry analysis using these cell lines and soluble ACE2 protein to confirm the expression and cell-surface display of the S-proteins (Supplementary data S4). Following this, we performed titrations using the biotinylated h-deface2 and p-deface2 to determine the ideal concentration of the proteins for the subsequent flow-cytometry experiments. We observed an initial increase in binding, as measured by Geometric Mean (Geo Mean FL-2-PE) of Flow-cytometry histograms for both h-deface2 and p-deface2 with the increase in the concentration of the proteins.

Interestingly, at higher concentrations, h-deface2 and p-deface2 displayed loss of binding to the cells, as evident from the “bell-shaped curve” (Supplementary data S5), suggesting that oligomerization of the proteins at high concentrations might be responsible for removing the engineered proteins from the solution phase. Therefore, based on the results from titration experiments, we selected the protein concentration of 0.1 μg/μL for both h-deface2 and p-deface2 to perform our subsequent assay to test the specificity of the proteins. We observed higher binding of both the h-deface2 and p-deface2 proteins to the Spike protein (Wuhan and Delta) expressing HEK293 cells than the control HEK293 cells (Supplementary data S6, S7, and Figure 6). However, a noticeable background binding of h-deface2 and p-deface2 to the control, HEK293 cells was also observed. Defensins are highly basic proteins, and the interaction of their positively charged amino-acid residues with negatively charged phosphatidyl-lipid moieties on the cell plasma membrane is most likely responsible for this background binding.

**Figure 6:**
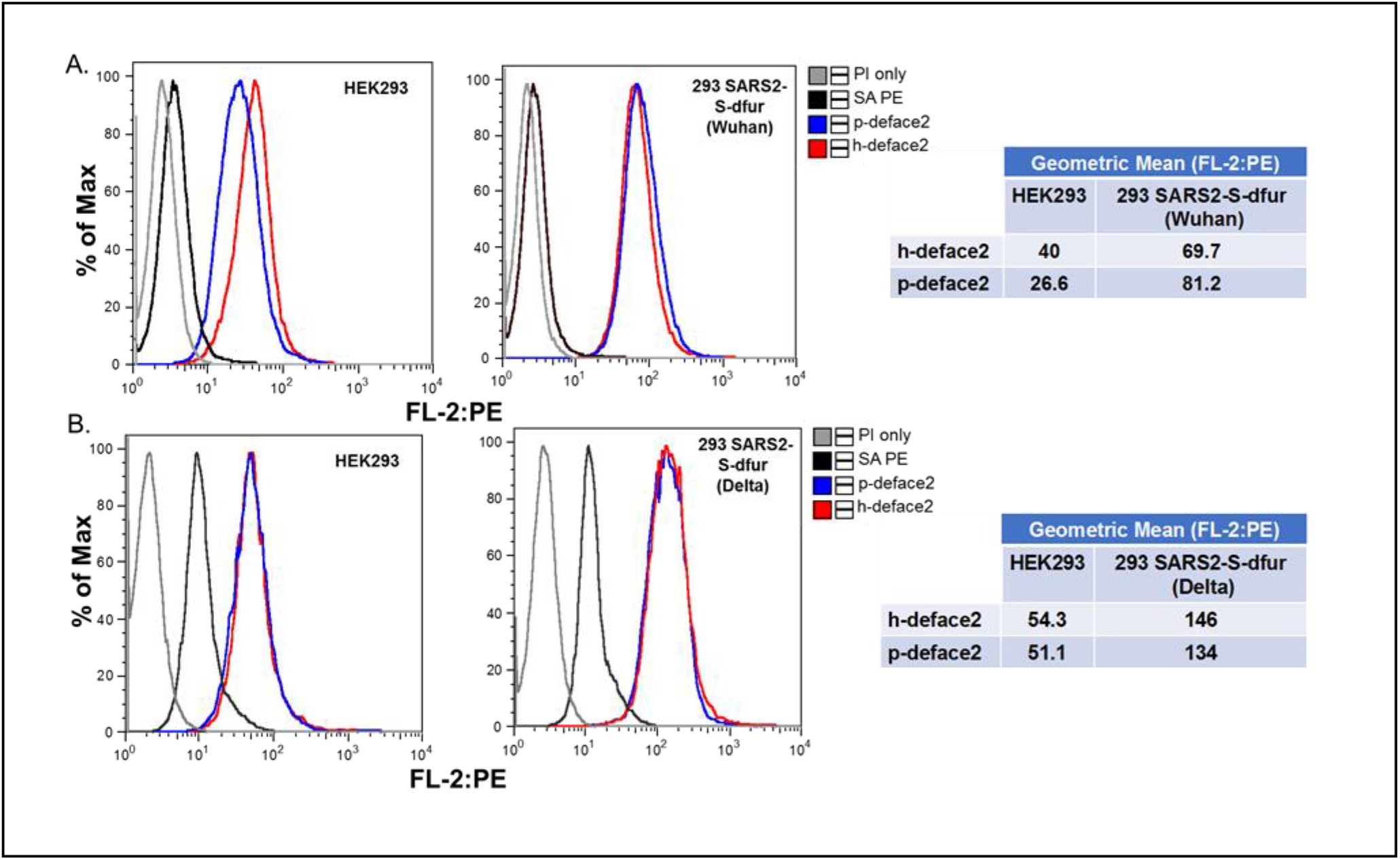
Flow-cytometry analysis of ACE2-defensin peptidomimetics binding to cells expressing SARS-CoV2 Spike-protein; HEK293 cell-lines expressing A) Wuhan-Hu-1 (D614) SARS-CoV2, and B) Delta (B.1.617.2) variant SARS-CoV2 Spike protein were incubated with h-deface2 or p-deface2 biotinylated proteins (0.1 μg/μL) conjugated to Streptavidin-Phycoerythrin (SA-PE) fluorophore. The Histogram overlay and the Geometric mean (FL-2:PE) data of the Flow-cytometry analysis are presented. Parental HEK293 cell-line were treated similarly with SA-PE conjugated h-deface2 or p-deface2 biotinylated proteins (0.1 μg/μL). SA-PE was used as secondary fluorophore control. All the cells were stained with Propidium Iodide (PI) for live-dead cell staining.

## Discussion

Early studies revealed that soluble ACE2 inhibits SARS-CoV-2 infection, showing that blocking the PPIs between ACE2 and Spike RBD might be a promising strategy for COVID-19 diagnosis and treatment (6). Since then, many trials using soluble ACE2 variants have been underway. However, although soluble ACE2 is a promising approach, it will be both cost and time prohibitive to mass-produce the protein in a eukaryotic expression system for deploying it in the production of COVID19 diagnostics or therapeutics. In this respect, peptide-based therapeutics present advantages, including cost and time efficiency in synthesis compared to eukaryotic proteins. Therefore, researchers are focusing on developing minimalist and efficient α-helical mimetic structures suitable for large-scale and cost-effective production (27). Most rationally designed peptidomimetics are based on ACE2 N-terminal helix 1 since this is the secondary structure that encloses most RBD interactions.

Unconstrained α-helical peptide inhibitors based on ACE2-helix 1 were able to block the interaction between Spike protein and ACE2 (27) with binding affinities at the millimolar range. Linear α-helices are flexible and less structured, frequently leading to lower affinity binders. Therefore, conformationally restricted ACE2 helix1-stapled peptides were produced, but affinity increased only to the micromolar range (28). Other stapled peptides were created and presented no evidence of viral neutralization (11), indicating that enhanced binding interactions are required to outcompete all the interactions of membrane-bound ACE2 and the virus Spike-protein to prevent viral infection effectively. In this regard, SARS-CoV-2 mini protein inhibitors containing residues 23 to 46 from helix 1 were also developed presenting dissociation constant at a low nanomolar range (29). Mini proteins are a diverse group of protein scaffolds characterized by small (1–10 kDa) size, stability, and versatility (30, 31), being currently used as scaffolds for protein engineering. These SARS-CoV-2 mini protein inhibitors were computer-generated and effectively targeted the ACE2-RBD interface, however, as non-natural scaffolds, little is known about their stability or cellular toxicity (29).

Expanding the portfolio of nontoxic, stable, and inexpensive ACE2 peptidomimetics is crucial for producing SARS-CoV-2 detection probes, detecting neutralizing antibodies, and generating new therapeutic strategies. Therefore, our study intended to create conformationally constrained α-helices structured on a prokaryotically expressed and highly stable tertiary structure to generate potent and stable binders of SARS-CoV-2 Spike RBD. We used defensins as scaffolds, which are small and highly stable proteins found in nature. Defensins are efficiently expressed in *E. coli*, nontoxic for eukaryotes, present exceptional thermostability, and often enclose a disulfide-stabilized α-helix. We reasoned that defensins would provide a well-structured framework to display critical ACE-2 residues on a constrained α-helical structure.

Defensins exhibit antimicrobial properties against pathogenic organisms, including bacteria, fungi and viruses (32). Recently, a human beta-defensin expressed in the human respiratory epithelium was shown to bind to the Spike RDB and inhibit SARS-CoV-2 pseudovirus infection on ACE2 expressing HEK 293Tcells (33). Earlier studies also demonstrated the binding of a fungal defensin into the Spike RBD from SARS-CoV-2 (34). However, unlike our study, in both these cases, the defensin alpha-helix was not engineered to mimic the ACE2 alpha helix1, and perhaps for that reason, their binding affinities for SARS-CoV-2 Spike protein were observed at micromolar ranges.

The engineered proteins were highly expressed in a bacterial expression system, and purified to homogeneity, demonstrating exceptional thermostability and potent binding against SARS-CoV-2 Spike protein. We observed that the apparent dissociation constant for h-deface2 is higher than p-deface2. The reason is likely due to the shortened length of the human defensin α-helix. The use of plant defensin provided a more extended α-helix that accommodated ACE2 residues 29 to 39, improving binding as observed by a lower apparent *K*_d_ value for p-deface2. Based on the three-dimensional structure reported earlier, we noticed that ACE2 displays a palindromic region where Histidine 34 is centered and the pairs L29/D30 and D38/L39 are equidistant located from H34. We reasoned that this palindromic positioning could allow spike docking in both orientations, and therefore we introduced these additional residues into p-deface2.

Furthermore, the addition of K31N and E35K mutations in p-deface2-MUT culminated with improved binding (*K*_d_ 14.4 nM). Therefore, the defensin structure was not only able to incorporate the function of the native ACE2 residues for binding, but remarkably it was also able to achieve a gain of function with the same specific mutations discovered in the evolution of ACE2 derivatives (26), demonstrating that they are versatile scaffolds. *K*_d_ calculations using the BLI assay were not possible since curves did not follow a single exponential behavior, indicating that multiple binding events were likely occurring. Defensins tend to dimerize through their β-sheet face (35). Additionally, the SARS-CoV Spike glycoprotein undergoes oligomerization (36). It is possible that binding to multimeric structured defensins and Spike protein produces a more complex interaction that did not allow proper kinetic analysis through BLI assays.

In addition, the engineered proteins distinguished HEK293/SARS-CoV-2 cell lines from the parental HEK293 control cells at the conditions tested. However, a high background binding was observed, which is likely due to the defensins mode of action that involves an electrostatic interaction with the pathogen cell membrane due to the highly cationic character of defensins. Studies have demonstrated that the binding of defensins to cell membrane phosphatidic acids is part of their mechanism of action (37, 38). Although it might sound contradictory, cell surface stickiness might be a wanted feature of such scaffolds since their local use into the respiratory tract may be favored by a diminished local clearance. Mutations of the engineered proteins are being studied to evaluate this hypothesis by our group.

In addition to the observed thermostability, the proteins retained total biological activity upon lyophilization and reconstitution, important features considering their potential use as point of care diagnostic reagents.

The initial focus of our study was directed against the SARS-CoV-2 that emerged in 2019, but the current emergence of genetic variants introduces new challenges that need to be addressed. Furthermore, recent data demonstrated that therapeutic intervention based on ACE2 hotspot residues is a promising strategy to hit emerging variants and similar SARS viruses (13). Accordingly, the engineered defensin-ACE2 peptides demonstrated binding towards the Wuhan and Delta variant SARS-CoV-2.

α-Helices play a significant role in mediating protein-protein interactions in nature, being stabilized via noncovalent interaction within proteins’ tertiary structures. Although chemical biologists frequently use synthetic strategies to stabilize the α-helical conformation of peptides, other strategies involve using mini-proteins as scaffolds to display stable α-helical domains (12). To our knowledge, defensins have not yet been explored as mini protein scaffolds, and our strategy demonstrated that they might represent a superior strategy for developing constrained alpha-helices mimetic and high-affinity binders for additional targets.

## Material and Methods

### Material

SARS-CoV-2 Spike protein (ECD) was purchased from Genescript, USA, Inc. Chemical reagents were purchased from Sigma Aldrich. Biolisa COVID-19 Anticorpo Neutralizante kit (Ref: K243-1) was kindly provided by Bioclin^®^.

### Cell Lines overexpressing SARS-CoV2 Spike protein

HEK293 cells overexpressing 1) Wuhan-Hu-1 (D614) (Catalogue code: 293-cov2-sdf) or 2) Delta/B.1.617.2 variant) (Catalogue code: 293-SARS2-S-V8-dfur) were purchased from Invivogen. The cells were grown in DMEM media containing 10% Fetal Bovine Serum, and 1% penicillin-streptomycin antibiotic following the standard tissue culture method. The parental HEK293 cells were used as a control for binding specificity.

### Rational design of defensin-based peptidomimetics

The ACE2-RBD interface was analyzed using the Protein Data Bank (PDB) annotated structure 2AJF. Analysis of the complex structure reveals that most RBD contacts involve the surface located ACE2 helix 1 residues L29, D30, K31, N33, H34, E35, E37, D38, and L39. Therefore, we positioned the ACE2 Histidine 34 at the center of the human defensin α-helix on its most solvent-accessible face. Additional residues D30, K31, and E35, were modeled into the helix to produce h-deface2.

Likewise, p-deface2 was engineered by positioning H34 from ACE2 at the center of the plant defensin α-helix, then modeling the additional residues (L29, D30, K31, N33, E35, E37, D38, and L39) at the flanking regions of defensin helix, to generate p-deface2. Finally, the hypothetical three-dimensional structure of defensin-based peptides were created using The Phyre2 web portal for protein modeling, prediction, and analysis (39). All genes were purchased from Biomatik Corporation (Ontario, CA) and cloned into pET-32a (+) between the *Nco* I and *Xho* I restriction sites. Therefore, all proteins contained a Trx-Tag, followed by a His-tag and an S-tag (Supplementary data S1). Trx-tag was included in the construct to assist disulfide bond formation. Control experiments using proteins lacking Trx-Tag were performed and indicated accordingly (Supplementary data S2).

### Protein Expression and Purification

Recombinant plasmids were transformed by electroporation into *Escherichia coli* pLysS (DE3) cells. The transformed cells were plated on LB agar plates containing 100μg/mL of ampicillin and 37.5 μg/mL chloramphenicol, and isolated colonies were inoculated in LB liquid medium containing the same antibiotics and cultured overnight at 37 °C until reaching the OD at the 600 nm wavelength of 0.6 absorbances. The cells were induced with 0.1 mM IPTG and incubated at 37 °C for 7 h. For purification, all procedures were performed at 4 °C. The cells were suspended in 20 mL of 50 mM Tris-HCl, pH 8.0: 500 mM NaCl (Buffer A) containing 1mM phenylmethylsulfonyl fluoride (PMSF) and lysozyme (10 mg/mL). Cells were disrupted by sonication on ice for 30 min. Cell debris were removed by centrifugation at 10,000 g for 10 minutes, and the supernatant was applied to a Buffer A pre-equilibrated Ni-NTA Sepharose column. The weakly bound proteins were removed by washing with buffer B (Tris-HCl 50 mM pH 8.0; 500 mM NaCl; 60 mM imidazole), and proteins were eluted in buffer C (500 mM NaCl, 500 mM Imidazole; 50 mM Tris-HCl, pH 8.0). The eluted was dialyzed against PBS pH 7.2 (Dulbecco’s Phosphate Buffered Saline SIGMA), mixed to 50% glycerol, and stored at-20°C. SDS-PAGE determined the purity.

Biotinylated proteins were obtained using Biotin N-hydroxysuccinimide ester reagent (Sigma Aldrich) according to the manufacturer’s protocol.

### Lyophilization and Reconstitution

Purified Defensin-ACE2 peptidomimetics were stored in PBS with 4% mannitol (w/v). Samples were frozen at −80°C for 12 hours and dried for 24 hours using a workbench freeze dyer (L101-Liobras) with a condenser temperature presented at −51°C and vacuum < 55 Hg. The dried samples were tightly capped and stored at 4°C. The lyophilized samples were reconstituted by adding the appropriate amount of ultrapure water. Glycerol was added to 50% final concentration and samples were stored at −20°C.

### Biolayer interferometry (BLI)

BLI was performed on an Octet Red96 at 25 °C. Binding reactions were performed in 1X kinetic buffer (Sartorius, 18-1105) consisting of 20 mM phosphate buffer, pH 7.6, 2mM KCl, 150mM NaCl, 0.02% Tween 20, 0.1% BSA, and 0.05% sodium azide. Biotinylated defensin-based peptidomimetics were immobilized on streptavidin-conjugated (SA) Biosensors (Sartorius, 18-5019) by dipping the sensor in 150nM solution until the signal was saturated (unless otherwise noted in the data). For nonspecific background binding of the S-receptor protein to the SA surface, a negative control sensor for every concentration tested was used to subtract individual background signal intensities. Kinetic experiments were performed separately for proteins to SARS-CoV-2 S-Protein. Experiments were performed using a gradient of concentrations and vary from experiment to experiment. Kinetic analysis was performed using Octet Data Analysis HT Software Version 12, with either a 1:1 or a 2:1 binding model. All Octet BLI experiments were performed at the Scripps Research Biophysics and Biochemistry Core.

### Enzyme-linked immunosorbent assay (ELISA) analysis

All ELISA assays were performed into 96 wells high binding microtiter plates. For direct ELISA assays, wells were coated (18 h, 4°C) with 25/50 ng of SARS-CoV-2 Spike ECD (Genescript) in 50 μL PBS (Phosphate buffer saline) and blocked with 200 μL of 1% BSA in TBST (PBS, 0,05% Tween-20) for 2h at 37°C. Next, biotin-coupled Defensins were diluted in TBST, 1% BSA, and added to the wells to a final volume of 100 μL. Finally, one hundred microliters of streptavidin-HRP (1:5000) was added to the wells and incubated for 15 minutes at room temperature. Plates were washed five times with PBST after each step, except for final washing, which included eight washing steps. The reaction was visualized by adding 50 μL chromogenic substrate (ONE STEP TMB, Scienco Biotech) for 15 min. The reaction was quenched with 50 μL Stop Solution, and absorbance at 450 nm was measured using an ELISA plate reader.

### Thermostability assay

Defensin-based ACE-2 peptidomimetics (1μg) were pre-incubated at 37°C, 50°C, 65°C, 80°C or 95°C for five minutes, cooled on ice for additional 5 minutes and mixed to 90 μL of PBST containing 1% of BSA and incubated for 1 hour against 50 ng of ECD previously immobilized on polystyrene ELISA plates. The wells were washed three times with PBST, followed by the addition of streptavidin-HRP (1:5000 dil), fifteen minutes incubation at room temperature, additional PBST washing, and finally, color development by the presence of the HRP substrate TMB. The addition of 0.16 M H2SO4 quenched reactions.

### Flow-Cytometry

Cells were grown to 80% confluency in tissue culture flasks. The adherent cells were then detached from the plates following the standard cell-detachment protocol using 0.05% Trypsin-EDTA (ThermoFisher Scientific, Catalog number: 25300054). Following this, the cells were resuspended in Flow cytometry (FCM) buffer (PBS containing 2% FBS and 0.1% sodium azide), and the cell number was estimated using a Vi-CELL XR Cell Viability Analyzer (Beckman Coulter). For the FCM experiments, approximately one-million cells were incubated in the FCM buffer with biotinylated ACE2 peptidomimetics (h-deface2 and p-deface2) labeled with Streptavidin-phycoerythrin (SA PE, Prozyme) at the indicated protein concentrations (for one hour at 4°C in the dark) in a total reaction volume of 100 μL. Propidium Iodide (PI) (eBiosceince, Catalog number 00-6990-50) (2μL) was also added to the reaction to estimate cell viability. The cells were then washed twice with 500 μL FCM buffer, and the cell pellets were resuspended back in 500 μL of FCM buffer for analysis in Becton-Dickinson FACSCalibur. The data were analyzed using FlowJo software (BD Biosciences). For FCM assay using hACE2-Fc (Invivogen, Catalogue code: fc-hace2), 5μL of the protein was incubated for 1 hour at 4°C with the cells using a similar assay condition described above. The cells were washed once (500 μL FCM buffer) and resuspended in 100 μL of the FCM buffer with 5μL of the secondary Antibody (PE-anti human IgG Fc Recombinant, Biolegend, Catalogue 366903) and incubated for 30 minutes at 4°C., in the dark. The cells were finally washed as described before and resuspended in 500 μL FCM buffer for Flow-analysis. PI was used for the estimation of cell viability

## Abbreviations

(COVID-19): Coronavirus Disease 2019
(SARS-CoV-2): severe acute respiratory syndrome coronavirus 2
(RBD): receptor binding domain
(ECD): extracellular domain
(ACE2): Angiotensin Converting Enzyme 2

## Supplementary data

**Supplementary Figure S1:** Schematic diagram of Defensin-ACE2 constructs.

**Supplementary Figure S2:** Binding of Defensin-ACE2 proteins to SARS-C2V-2 Spike protein in the presence and absence of Trx-tag.

**Supplementary Figure S3:** Effect of lyophilization and reconstitution on deface2 ability to detect SARS-CoV-2 spike protein.

**Supplementary Figure S4:** Soluble ACE2 binds SARS-CoV2 Spike protein expressing HEK293 cells.

**Supplementary Figure S5:** Binding titrations using ACE2-defensin peptidomimetics to HEK293 SARS-CoV2 Spike expressing cells.

**Supplementary Figure S6:** Flow-cytometry analysis of ACE2-defensin peptidomimetics binding to cells expressing Wuhan-Hu-1 (D614) SARS-CoV2 Spike protein.

**Supplementary Figure S7:** Flow-cytometry analysis of ACE2-defensin peptidomimetics binding to cells expressing Delta (B.1.617.2) SARS-CoV2 Spike protein.

## Acknowledgments

This work received financial support from SENAI Green Chemistry-ABDI inside the SGF project n° 329254 and Grant PAP UDESC FAPESC 2021TR889. We would like to thank Dennis Young of Flow-Cytometry Shared Resource (UC San Diego Moores Cancer Center) for help with Flow-cytometry, Nicholas Lee and J.A. Hammond at the Scripps Research Biophysics Biochemistry Core for the Octet BLI experiments. We want to also thank Bioclin^®^ for kindly providing the Biolisa COVID-19 Anticorpo Neutralizante kit (Ref: K243-1).

## Conflict of interest

The authors intend to file a provisional patent application regarding the engineered proteins.

