## SupplementaryMaterial for "Engineering Defensin α-helix to produce high-affinity SARS-CoV-2 Spike protein binding ligands"

**Supplementary Material**

**Supplementary Figure S1: Schematic diagram of Defensin-ACE2 constructs. Panel A: Schematic diagram of h-deface2 and p-deface2 primary structure.** Trx-Tag, Histidine-tag and S- tag are highlighted in gray, cyan and green, respectively. h-deface and p-deface amino acid sequences are highlighted in orange. ACE2 inserted residues are shown in red. Cysteine involved in disulfide bridges are indicated with an asterisk**. Panel B: Predicted Three-dimensional structure of h-deface2 and pdeface2.** The hypothetical three-dimensional structure of defensin-based peptides were created using The Phyre2 web portal for protein modeling, prediction, and analysis. β-sheets are represented in yellow, α-helices are represented in blue and helix-stabilizing disulfides are shown in red.

**Supplementary Figure S2: Binding of Defensin-ACE2 proteins to SARS-C2V-2 Spike protein in the presence and absence of Trx-tag.** Elisa assays were performed using 50 ng immobilized SARS-CoV-2 Spike protein and biotinylated defensin-ACE mimetic in the presence and absence of Trx-tag.

**Supplementary Figure S3: Effect of liophilization and reconstitution on deface2 ability to detect SARS-CoV-2 spike protein.** Elisa assays were performed using 50 ng immobilized SARS-CoV-2 Spike protein and biotinylated defensin-ACE mimetic before and after lyophilization.

**Supplementary Figure S4: Soluble ACE2 binds SARS-CoV2 Spike protein expressing HEK293 cells.** Histogram overlay plots of the Flow-cytometry performed on HEK293 cell-lines expressing either A) Wuhan-Hu-1 (D614) or B) Delta variant (B.1.617.2) SARS-CoV2 Spike proteins using soluble hACE2-Fc and secondary-antibody (PE anti-human IgG-Fc) (Red) or only the PE anti-human IgG-Fc antibody (Blue) as the secondary-antibody control. The parental HEK293 cells were used as control cell line. All cells were also stained with Propidium Iodide (PI) for live-dead cell staining.

**Supplementary Figure S5: Binding titrations using ACE2-defensin peptidomimetics to HEK293 SARS-CoV2 Spike expressing cells.** HEK293 cell-lines expressing Wuhan-Hu-1 (D614) SARS-CoV2 Spike proteins were incubated with either A) h-deface2 or B) p-deface2 biotinylated proteins conjugated to Streptavidin-Phycoerythrin (SA-PE) fluorophore at the indicated concentrations. Histogram overlay plots (upper panel) and the Geometric Mean (FL-2:PE) plotted against the protein concentrations (lower panel) from the Flow-cytometry measurements are presented. SA-PE was used as secondary fluorophore control. All the cells were stained with Propidium Iodide (PI) for live-dead cell staining.

**Supplementary Figure S6: Flow-cytometry analysis of ACE2-defensin peptidomimetics binding to cells expressing Wuhan-Hu-1 (D614) SARS-CoV2 Spike protein**. HEK293 cell-lines expressing Wuhan-Hu-1 (D614) SARS-CoV2 Spike protein was incubated with either h-deface2 or p-deface2 biotinylated proteins (0.1 μg/μL) conjugated to Streptavidin-Phycoerythrin (SA-PE) fluorophore (upper panel). Parental HEK293 cell-line were also treated similarly with SA-PE conjugated h-deface2 or p-deface2 biotinylated proteins (0.1 μg/μL) to test the binding specificity of the engineered proteins (lower panel). Histograms, Geometric mean, and the dot-plots data of the flow-cytometry analysis are presented. SA-PE was used as secondary fluorophore control. All the cells were stained with Propidium Iodide (PI) for live-dead cell staining.

**Supplementary Figure S7: Flow-cytometry analysis of ACE2-defensin peptidomimetics binding to cells expressing Delta (B.1.617.2) SARS-CoV2 Spike protein**. HEK293 cell-lines expressing Delta (B.1.617.2) variant SARS-CoV2 Spike protein was incubated with either h-deface2 or p-deface2 biotinylated proteins (0.1 μg/μL) conjugated to Streptavidin-Phycoerythrin (SA-PE) fluorophore (upper panel). Parental HEK293 cell-line were also treated similarly with SA-PE conjugated h-deface2 or p-deface2 biotinylated proteins (0.1 μg/μL) to test the binding specificity of the engineered proteins (lower panel). Histograms, Geometric mean, and the dot-plots data of the flow-cytometry analysis are presented. SA-PE was used as secondary fluorophore control. All the cells were stained with Propidium Iodide (PI) for live-dead cell staining.

Figure S1

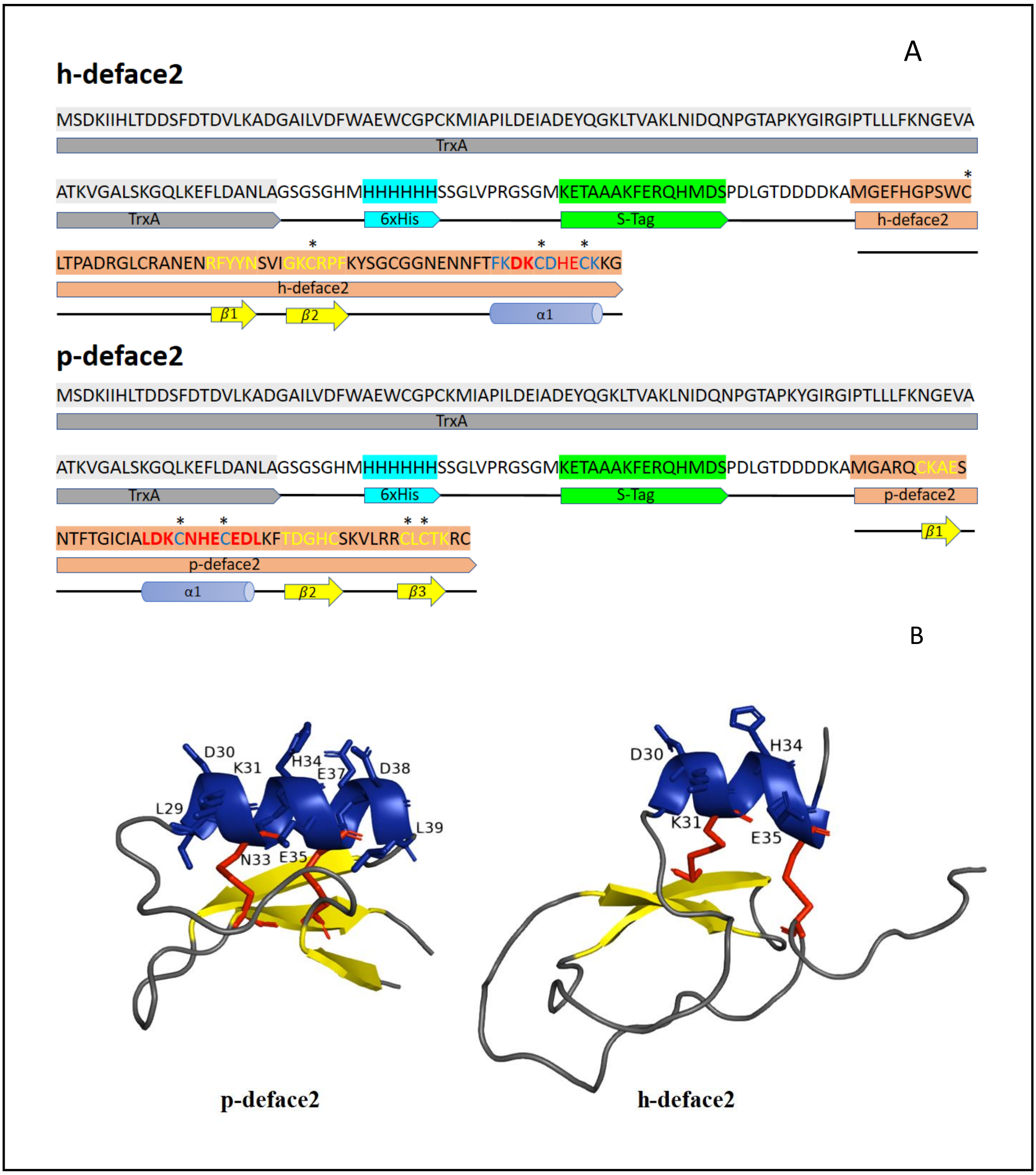
A
h-deface2
MSDKIIHLTDDSFDTDVLKADGAILVDFWAEWCGPCKMIAPILDEIADEYQGKLTVAKLNIDQNPGTAPKYGIRGIPTLLLFKNGEVA
TrxA
*
ATKVGALSKGQLKEFLDANLAGSGSGHMHHHHHHSSGLVPRGSGMKETAAAKFERQHMDSPDLGTDDDDKAMGEFHGPSWC
TrxA
6xHis
S-Tag
h-deface2
* * *
LTPADRGLCRANENRFYYNSVIGKCRPFKYSGCGGNENNFTFKDKCDHECKKG
h-deface2
β1
β2
α1
p-deface2
MSDKIIHLTDDSFDTDVLKADGAILVDFWAEWCGPCKMIAPILDEIADEYQGKLTVAKLNIDQNPGTAPKYGIRGIPTLLLFKNGEVA
TrxA
ATKVGALSKGQLKEFLDANLAGSGSGHMHHHHHHSSGLVPRGSGMKETAAAKFERQHMDSPDLGTDDDDKAMGARQCKAES
TrxA
6xHis
S-Tag
p-deface2
β1
* * * *
NTFTGICIALDKCNHECEDLKFTDGHCSKVLRRCLCTKRC
p-deface2
α1
β2
β3
B
D30
K31
H34
E37
D38
L29
L39
N33
E35
D30
H34
K31
E35
p-deface2
h-deface2

Figure S2

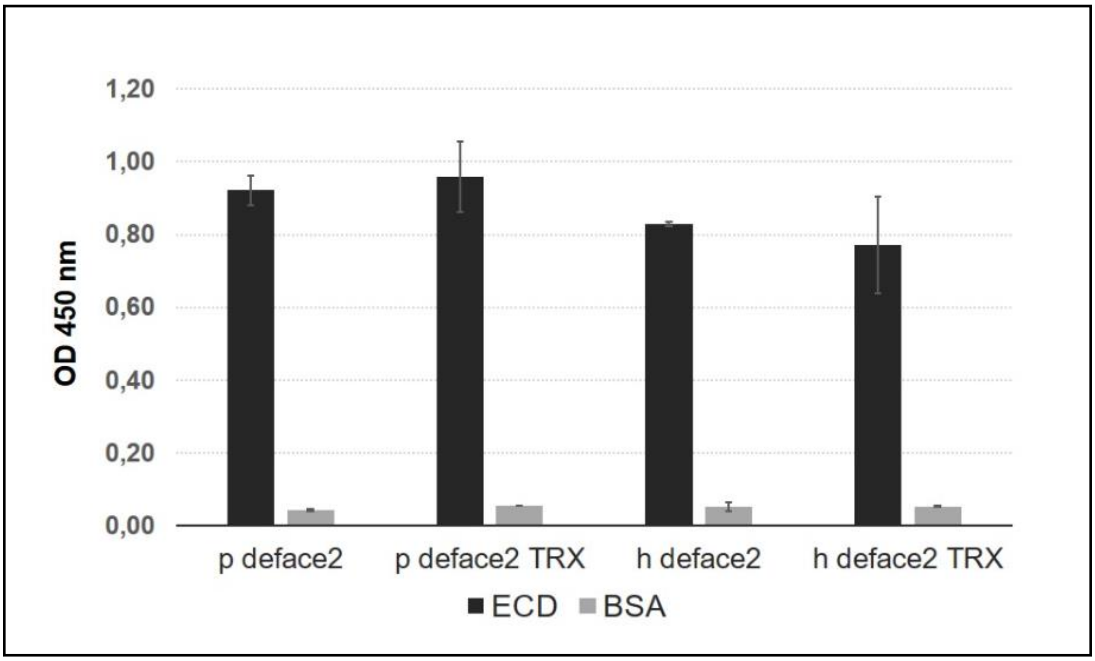
1,20
1,00
0,80
0,60
0,40
0,20
0,00
OD 450 nm
p deface2
p deface2 TRX
h deface2
h deface2 TRX
ECD
BSA

Figure S3

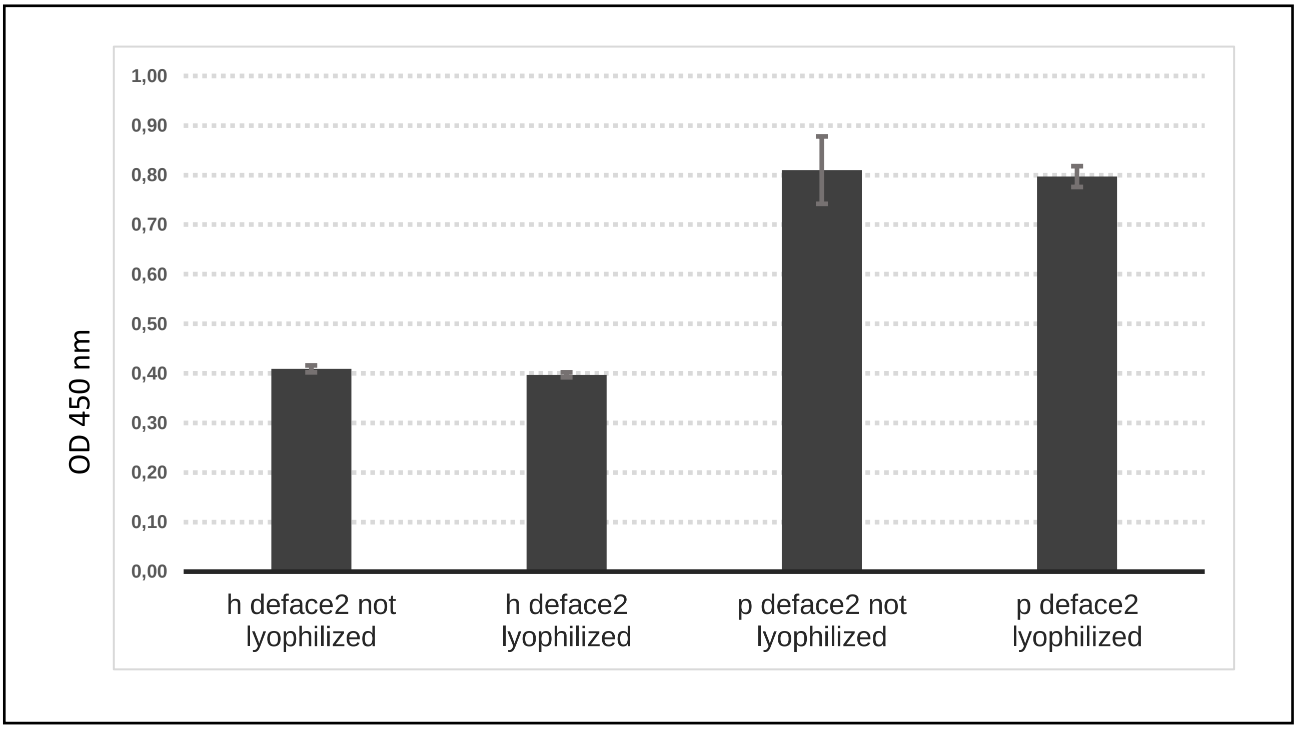
1,00
0,90
0,80
0,70
0,60
0,50
0,40
0,30
0,20
0,10
0,00
OD 450 nm
h deface2 not lyophilized
h deface2 lyophilized
p deface2 not lyophilized
p deface2 lyophilized

Figure S4

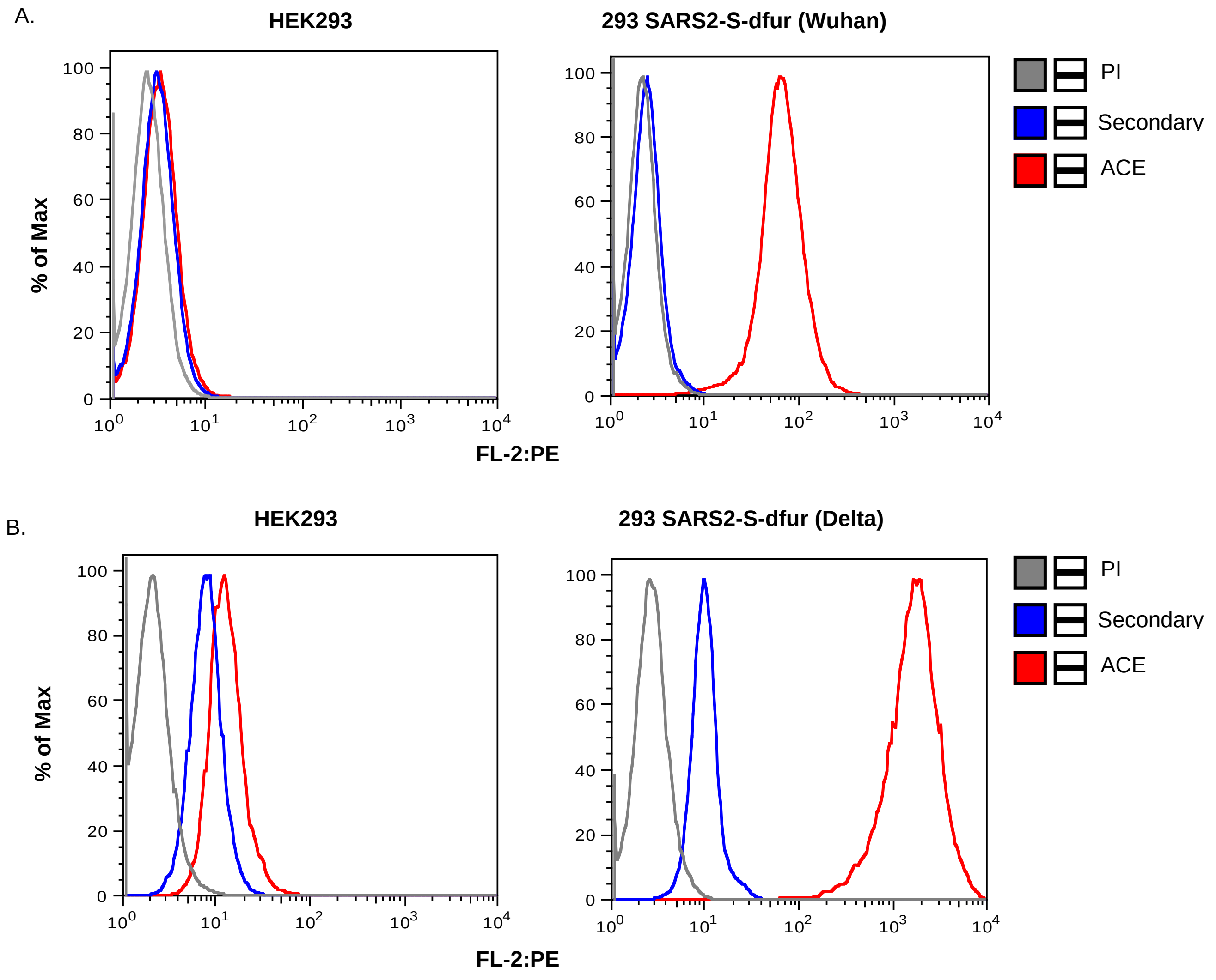
A.
HEK293
293 SARS2-S-dfur (Wuhan)
% of Max
FL-2:PE
PI
Secondary
ACE
B.
HEK293
293 SARS2-S-dfur (Delta)
% of Max
FL-2:PE
PI
Secondary
ACE

Figure S5

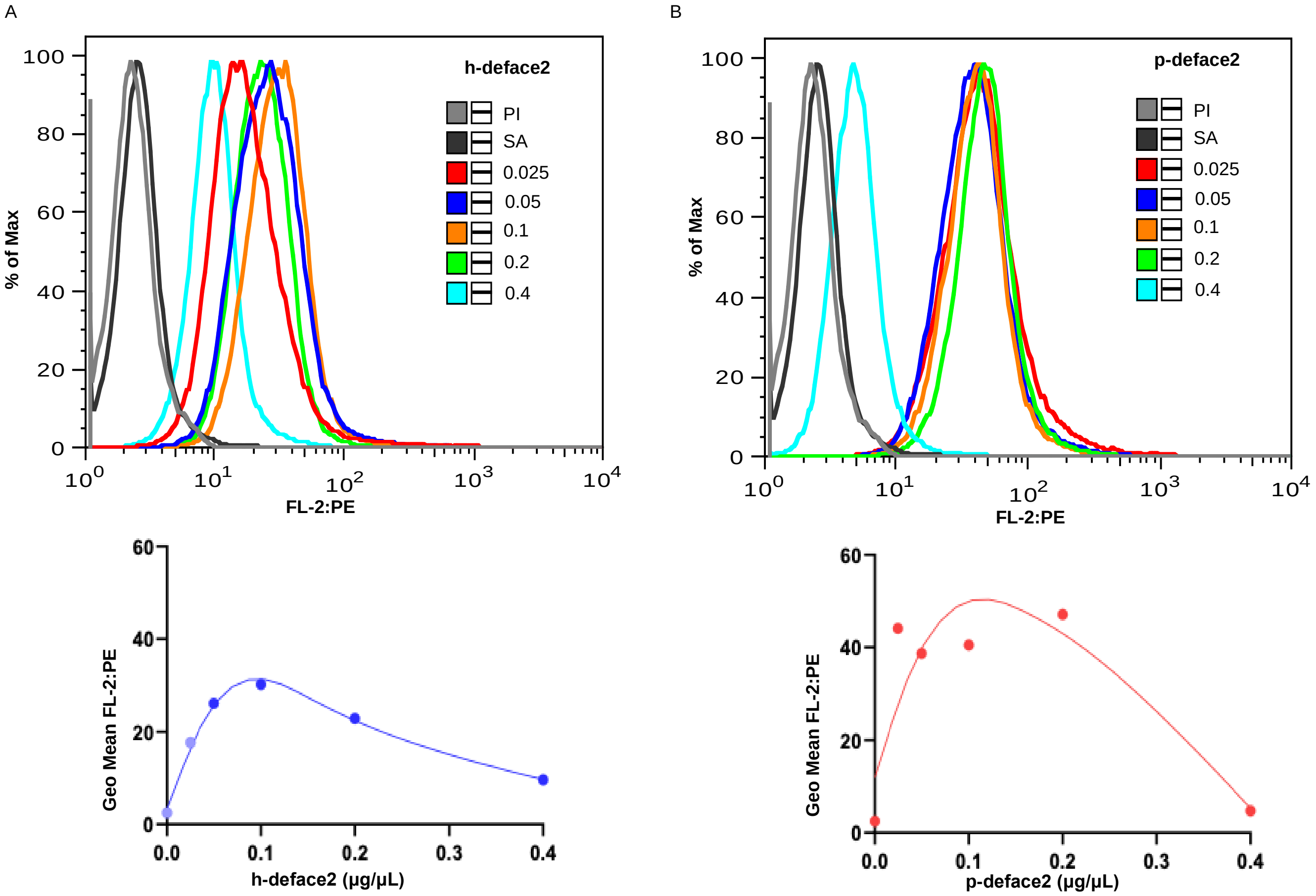
A
B
h-deface2
p-deface2
PI
SA
0.025
0.05
0.1
0.2
0.4
% of Max
FL-2:PE
Geo Mean FL-2:PE
h-deface2 (μg/μL)
p-deface2 (μg/μL)

Figure S6

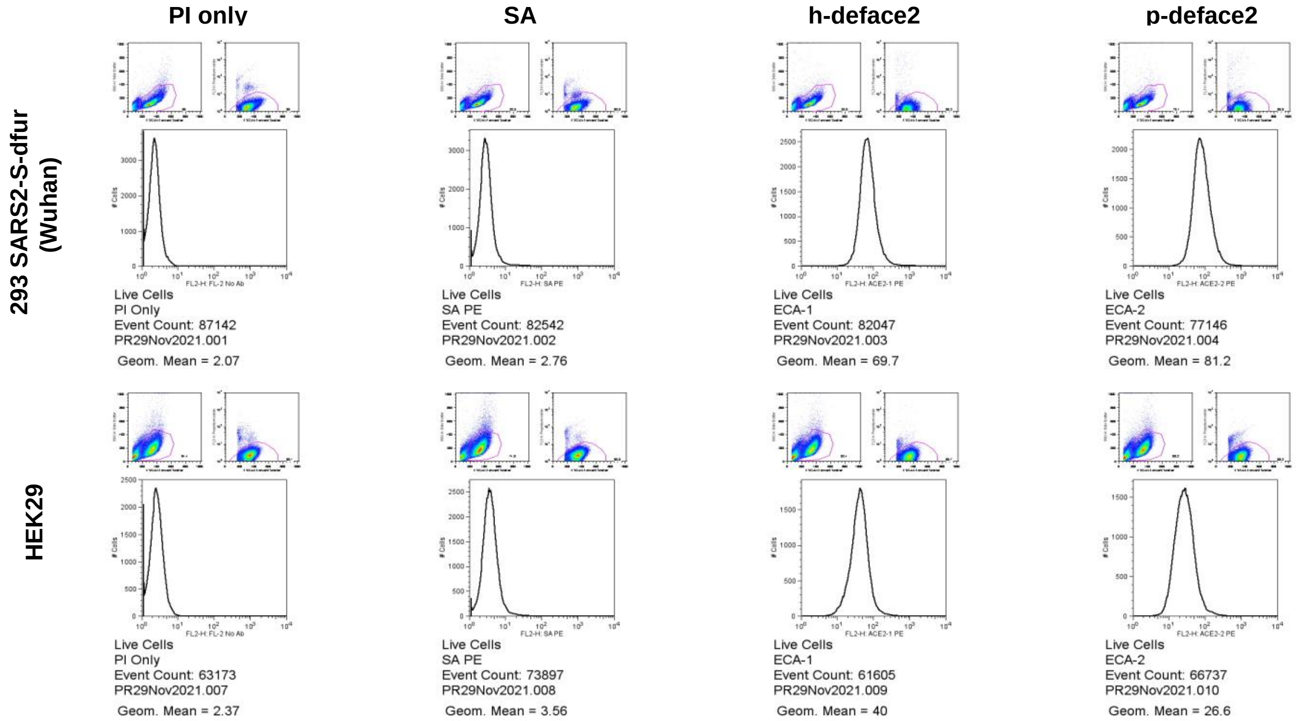

PI only
SA
h-deface2
p-deface2
293 SARS2-S-dfur (Wuhan)
HEK29
### Cells
FL2-H: FL-2 No Ab
FL2-H: SA PE
FL2-H: ACE2-1 PE
FL2-H: ACE2-2 PE
Live Cells
PI Only
Event Count: 87142
PR29Nov2021.001
Geom. Mean = 2.07
Live Cells
SA PE
Event Count: 82542
PR29Nov2021.002
Geom. Mean = 2.76
Live Cells
ECA-1
Event Count: 82047
PR29Nov2021.003
Geom. Mean = 69.7
Live Cells
ECA-2
Event Count: 77146
PR29Nov2021.004
Geom. Mean = 81.2
Live Cells
PI Only
Event Count: 63173
PR29Nov2021.007
Geom. Mean = 2.37
Live Cells
SA PE
Event Count: 73897
PR29Nov2021.008
Geom. Mean = 3.56
Live Cells
ECA-1
Event Count: 61605
PR29Nov2021.009
Geom. Mean = 40
Live Cells
ECA-2
Event Count: 66737
PR29Nov2021.010
Geom. Mean = 26.6

Figure S7

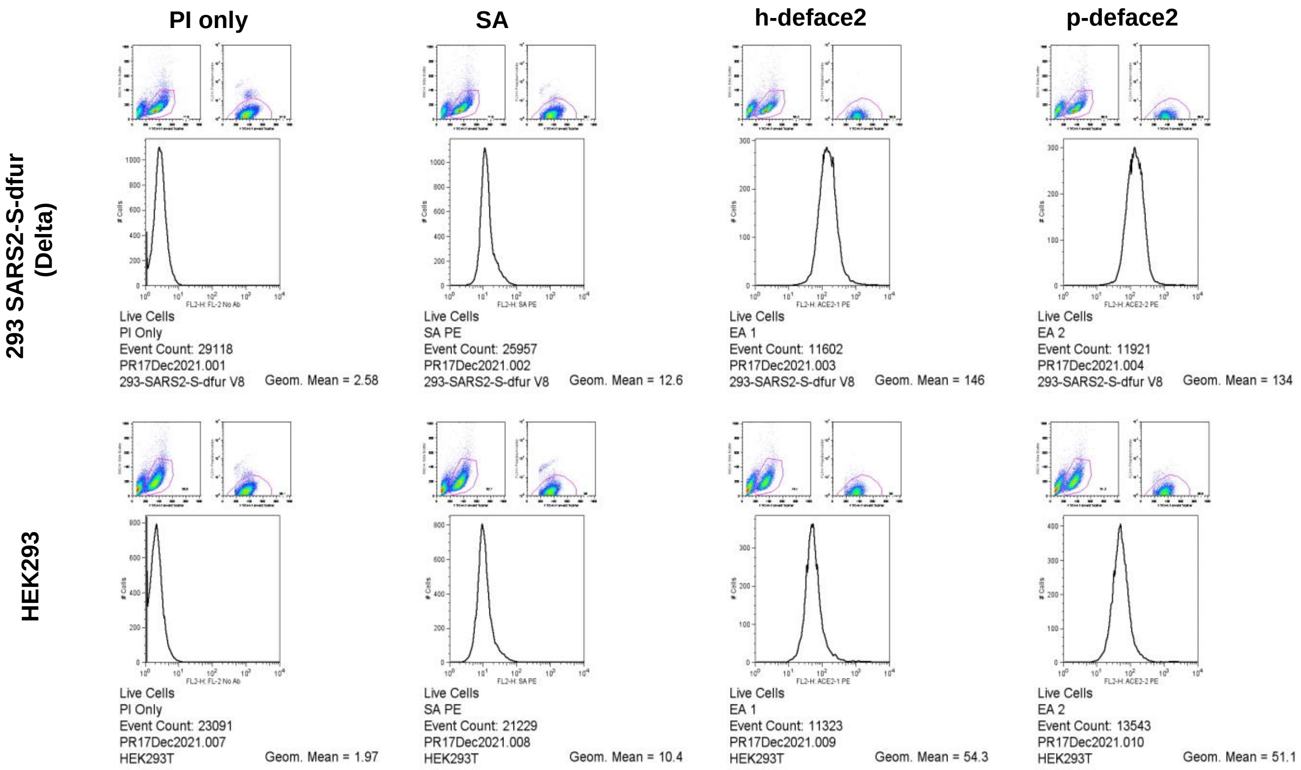
PI only
SA
h-deface2
p-deface2
293 SARS2-S-dfur (Delta)
HEK293
Live Cells
PI Only
Event Count: 29118
PR17Dec2021.001
293-SARS2-S-dfur V8
Geom. Mean = 2.58
Live Cells
SA PE
Event Count: 25957
PR17Dec2021.002
293-SARS2-S-dfur V8
Geom. Mean = 12.6
Live Cells
EA 1
Event Count: 11602
PR17Dec2021.003
293-SARS2-S-dfur V8
Geom. Mean = 146
Live Cells
EA 2
Event Count: 11921
PR17Dec2021.004
293-SARS2-S-dfur V8
Geom. Mean = 134
Live Cells
PI Only
Event Count: 23091
PR17Dec2021.007
HEK293T
Geom. Mean = 1.97
Live Cells
SA PE
Event Count: 21229
PR17Dec2021.008
HEK293T
Geom. Mean = 10.4
Live Cells
EA 1
Event Count: 11323
PR17Dec2021.009
HEK293T
Geom. Mean = 54.3
Live Cells
EA 2
Event Count: 13543
PR17Dec2021.010
HEK293T
Geom. Mean = 51.1
